# Intra- and Inter-host Evolution of Human Norovirus in Healthy Adults

**DOI:** 10.1101/2023.05.30.542907

**Authors:** Sasirekha Ramani, Sara J. Javornik Cregeen, Anil Surathu, Frederick H. Neill, Donna M. Muzny, Harsha Doddapaneni, Vipin K. Menon, Kristi L. Hoffman, Matthew C. Ross, Ginger Metcalf, Antone R. Opekun, David Y. Graham, Richard A. Gibbs, Joseph F. Petrosino, Mary K. Estes, Robert L. Atmar

## Abstract

**Background:** Human noroviruses are a leading cause of acute and sporadic gastroenteritis worldwide. The evolution of human noroviruses in immunocompromised persons has been evaluated in many studies. Much less is known about the evolutionary dynamics of human norovirus in healthy adults.

**Methods:** We used sequential samples collected from a controlled human infection study with GI.1/Norwalk/US/68 virus to evaluate intra- and inter-host evolution of a human norovirus in healthy adults. Up to 12 samples from day 1 to day 56 post-challenge were sequenced using a norovirus-specific capture probe method.

**Results:** Complete genomes were assembled, even in samples that were below the limit of detection of standard RT-qPCR assays, up to 28 days post-challenge. Analysis of 123 complete genomes showed changes in the GI.1 genome in all persons, but there were no conserved changes across all persons. Single nucleotide variants resulting in non-synonymous amino acid changes were observed in all proteins, with the capsid VP1 and nonstructural protein NS3 having the largest numbers of changes.

**Conclusions:** These data highlight the potential of a new capture-based sequencing approach to assemble human norovirus genomes with high sensitivity and demonstrate limited conserved immune pressure-driven evolution of GI.1 virus in healthy adults.

## INTRODUCTION

Human noroviruses are a leading cause of acute gastroenteritis worldwide. Each year, norovirus infections result in an estimated 684 million cases of acute gastroenteritis globally, leading to a total of $4.2 billion spent in direct health system costs and over $60 billion in societal costs [1, 2]. In the United States alone, 19–21 million cases of human norovirus gastroenteritis occur annually, resulting in 56,000–71,000 hospitalizations and over $10 billion in associated costs [3, 4]. There are currently no licensed vaccines or targeted antivirals for norovirus, although four candidate vaccines are in clinical trials and several others are in preclinical development [5]. Given the significant clinical and economic burden of norovirus illness, the introduction of interventions is likely to be highly impactful.

Noroviruses are single stranded, positive sense RNA viruses belonging to the family *Caliciviridae*. The ∼7.5kb RNA genome is organized into three open reading frames (ORFs) of which ORF1 encodes for six nonstructural proteins, ORF2 codes for the major structural capsid protein, VP1, and ORF3 codes for the minor structural protein, VP2 [6]. Noroviruses are classified into ten genogroups (GI to GX) of which five genogroups are known to cause human infections [7]. Each genogroup is further divided into genotypes and some genotypes are further divided into variants. The prototype strain, Norwalk virus is assigned to genogroup 1 and is designated as genotype 1 (GI.1). The epidemiology of human noroviruses is complex with periodic emergence of new genotypes and new variants, particularly for the globally dominant GII.4 genotype. Multiple factors are proposed to contribute to the norovirus diversity including antigenic drift due to herd immunity, recombination events at the ORF1/ORF2 junction leading to chimeric viruses with better fitness, and accumulation of point mutations during replication [8].

Infections in immunocompromised persons can lead to chronic infection and illness and are associated with higher genetic diversity and elevated rates of evolution compared to acute infections. As such, infections in this population were considered one of the sources for emergence of new variants of noroviruses. However, a recent mathematical model showed that the relative rarity of such patients limits their impact on the norovirus epidemiology [9]. Given that noroviruses can be detected in stool for over 30 days from the onset of infection in healthy persons [10], another possibility is that inter- and intra-host evolution in healthy persons contributes to the introduction of new strains in the community.

While there have been several studies on human norovirus evolution in immunocompromised persons [11-21], there is limited data from healthy adults and children [22-26]. We previously established a controlled human infection model (challenge study) with the prototype Norwalk virus to determine the 50% human infectious dose (HID_50_) in healthy susceptible adults [27]. In the current study, we used sequential samples from the challenge study to evaluate intra- and inter-host evaluation of Norwalk virus in healthy persons. We performed full-length genome sequencing to determine (1) if there are changes in the virus genome over time among previously infected healthy adults; (2) if single nucleotide variants that arise are conserved between persons; and (3) whether the observed changes provide new insights into the biology of norovirus evolution.

## METHODS

### Study design and samples

The overall study design for the controlled human infection study with Norwalk virus has been described previously [27]. The study was registered at ClinicalTrials.gov (NCT00138476). Briefly, a randomized, double-blind, placebo-controlled evaluation of different dosages of Norwalk virus (4800, 48, 4.8 or 0.48 RT-PCR units) was performed in healthy adults 18–50 years of age. Virus-containing fecal samples, as identified using quantitative real-time and immunomagnetic capture RT-PCR assays [10], were used in the current analysis. Up to 12 samples post-challenge per person were tested, including one sample collected every day for the first week after challenge and at days 10, 14, 21, 28 and 56, where available. A schematic representation of study samples indicating volunteer ID, challenge dose group, clinical outcome (yes/no for acute gastroenteritis) and GI.1 detection in stool is shown in **Figure 1**.

**Figure 1:**
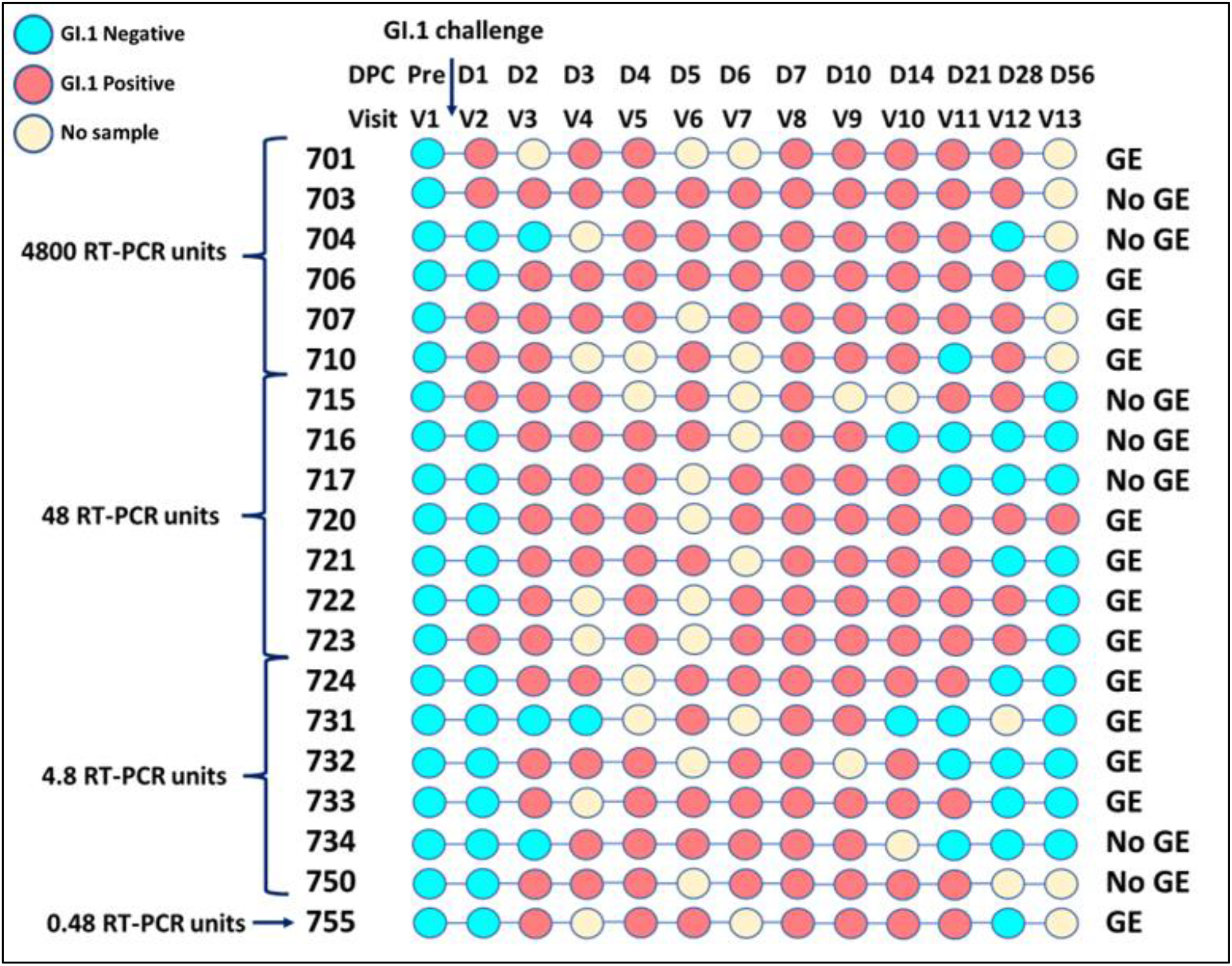
Schematic representation of the study samples. Study volunteer IDs (701-755), GI.1 challenge inoculum dose group, days/visits prior to- and post-challenge (DPC) are indicated. Persons who developed or did not develop gastroenteritis are marked as GE or no GE, respectively.

### RNA extraction, cDNA synthesis and library preparation

Stool samples were batched based on historical CT values for RNA extraction [10, 27]. Viral RNA was extracted using the Qiagen Viral RNA Mini Kit (QIAGEN Sciences, Maryland, USA) using the QIAcube automated extraction platform (QIAGEN, Hilden, Germany) according to the manufacturer instructions. RNA was analyzed using the RNA 6000 Nano assay (Agilent) or the RNA 6000 Pico assay for determination of RNA Integrity Number (RIN) and DV200 metrics. cDNA and strand-specific libraries were prepared utilizing New England Biolabs Inc. modules (E7525L and E7550L) without rRNA depletion or Poyy A+ RNA isolation steps as previously described [28]. Ilumina unique dual indices (20022370) were ligated to the individual cDNA products to barcode the samples and Kapa HiFi HotStart Library Amplification kit (kk2612, Roche Sequencing and Life Science) was used for library amplification.

### Norovirus-specific oligonucleotide capture-based sequencing

Eighty base pair length norovirus-specific capture probes were designed using 1,376 publicly available sequences resulting in a total of 39,300 unique probes (Twist Biosciences, Inc). The cDNA libraries were quantified, pooled and hybridized with biotin-labeled probes at 70°C for 16 hours [28]. Bound streptavidin beads were then washed and amplified with KAPA HiFi HotStart enzyme. cDNA libraries with Illumina adaptors were then batched into pools based on CT values and sequenced on Illumina NovaSeq S4 flow cells to generate 2×150 bp paired-end reads.

### Full-length norovirus genome assembly and data analysis

Quality trimmed (Phred score > 25) and host-filtered reads were processed through VirMAP for generating full-length norovirus genomes [29]. Briefly, VirMAP uses a combination of a mapping assembly algorithm combining nucleotide and amino-acid alignment information in a tiered manner to build virus-like super-scaffolds. This is followed by a merging system designed to hybridize the mapping assembly super-scaffolds and de-novo assembly contigs using an improvement algorithm that iteratively merges and rebuilds contig information. Finally, a taxonomic classification algorithm centered around a bits-per-base scoring system classifies the final reconstructions. Final reconstructions assembled using this process were manually inspected using Geneious Prime® 2022.1.1 and aligned against the reference genome to determine the quality of assemblies. A reconstructed genome with >90% the length of the Norwalk virus reference genome, NC_001959.2, was considered a complete genome. Genomes with above 90% completeness were further processed through a variant calling pipeline. Briefly, reads were aligned against the reference sequence using BWA and variant calling was done using iVar and then annotated with SnpEff. Positions with a minimum coverage of 20 reads and >10% minor variant frequency were plotted for analysis of single nucleotide variants. Further analysis was done using R.

## RESULTS

### Study samples and sequencing statistics

A total of 156 GI.1 positive samples were sequenced in the present study. The samples were collected from 13 persons with GI.1-associated gastroenteritis and 7 persons who did not meet the study’s clinical definition of gastroenteritis **(Table 1)** [10, 27]. Over 3 billion total raw reads were generated from the samples, with 79% percent of reads kept post quality trimming and host filtering. Very few reads (0.03%) mapped to the human genome as expected. A total of 24% of pre-processed reads mapped to the GI.1 reference genome.

**Table 1:**
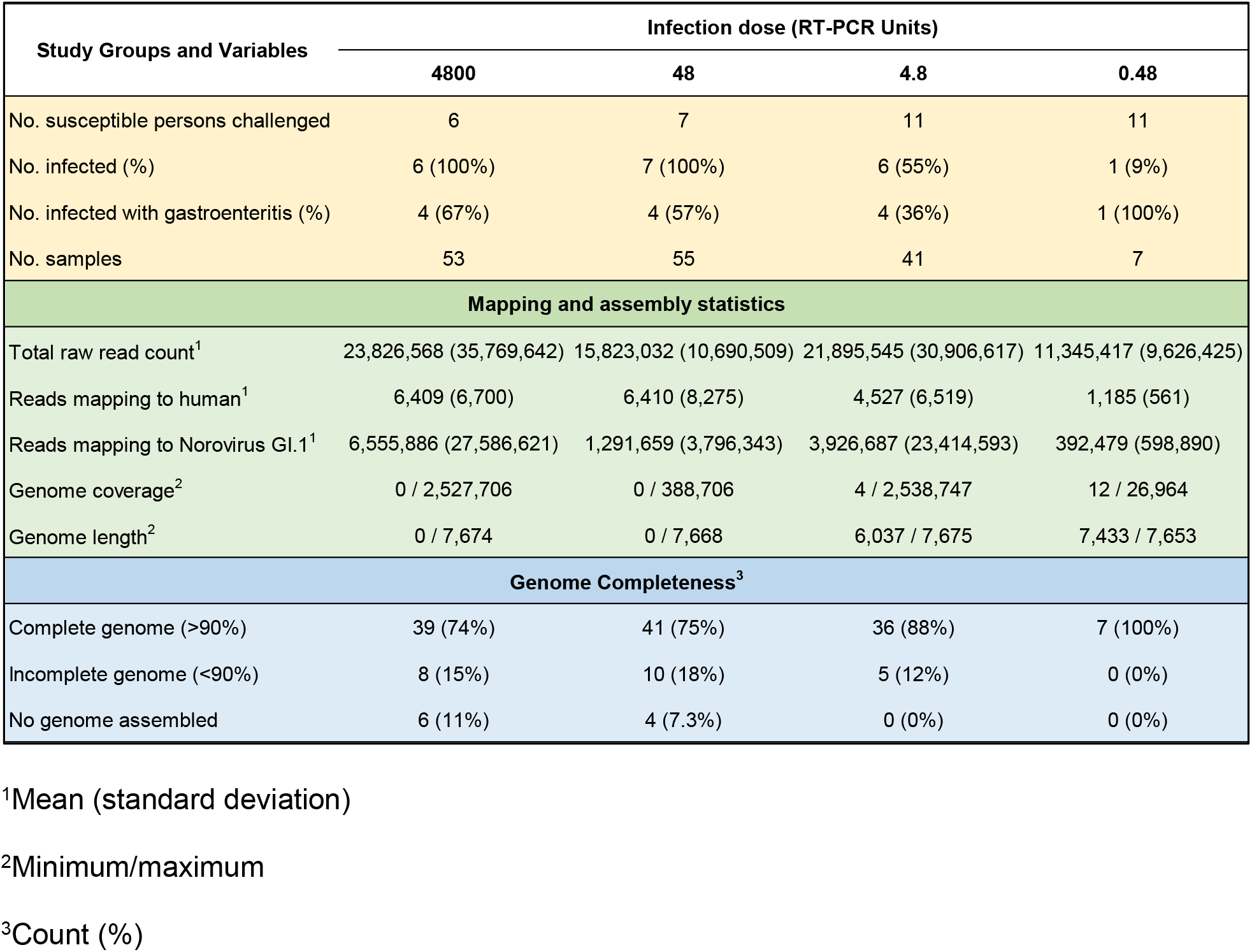
Study samples and sequencing read statistics for the different dose groups.

| Study Groups and Variables | Infection dose (RT-PCR Units) |  |  |  |
| --- | --- | --- | --- | --- |
|  | 4800 | 48 | 4.8 | 0.48 |
| No. susceptible persons challenged | 6 | 7 | 11 | 11 |
| No. infected (%) | 6 (100%) | 7 (100%) | 6 (55%) | 1 (9%) |
| No. infected with gastroenteritis (%) | 4 (67%) | 4 (57%) | 4 (36%) | 1 (100%) |
| No. samples | 53 | 55 | 41 | 7 |
| Mapping and assembly statistics |  |  |  |  |
| Total raw read count<sup>1</sup> | 23,826,568 (35,769,642) | 15,823,032 (10,690,509) | 21,895,545 (30,906,617) | 11,345,417 (9,626,425) |
| Reads mapping to human<sup>1</sup> | 6,409 (6,700) | 6,410 (8,275) | 4,527 (6,519) | 1,185 (561) |
| Reads mapping to Norovirus GI.1<sup>1</sup> | 6,555,886 (27,586,621) | 1,291,659 (3,796,343) | 3,926,687 (23,414,593) | 392,479 (598,890) |
| Genome coverage<sup>2</sup> | 0 / 2,527,706 | 0 / 388,706 | 4 / 2,538,747 | 12 / 26,964 |
| Genome length<sup>2</sup> | 0 / 7,674 | 0 / 7,668 | 6,037 / 7,675 | 7,433 / 7,653 |
| Genome Completeness<sup>3</sup> |  |  |  |  |
| Complete genome (>90%) | 39 (74%) | 41 (75%) | 36 (88%) | 7 (100%) |
| Incomplete genome (<90%) | 8 (15%) | 10 (18%) | 5 (12%) | 0 (0%) |
| No genome assembled | 6 (11%) | 4 (7.3%) | 0 (0%) | 0 (0%) |

<sup>1</sup>Mean (standard deviation)
<sup>2</sup>Minimum/maximum
<sup>3</sup>Count (%)

### GI.1 genomes can be assembled with high sensitivity using capture-based sequencing

Complete GI.1 genomes were generated in 123 of 156 samples (79%) while partial genomes could be assembled in an additional 23 samples (15%) **(Table 1)**. The number of samples with complete genomes in each dose group of the study and for volunteer are shown in **Figures 2 and 3**, respectively. With the exception of a day 6 post-challenge sample from volunteer 723, complete genomes were assembled in all samples (99%) with detectable CT values. Importantly, we could assemble complete genomes in 45% of samples with CT values below the limit of detection (< 36 cycles) of the GI.1 RT-qPCR used in our study. Complete genomes were assembled from samples collected as late as 28 days post-challenge. These data suggest that a probe capture-based sequencing method allows for complete assembly of human norovirus genomes with high sensitivity and that genome assembly was independent of challenge inoculum dose. We also identified a GII.4 (Hunter variant) strain in a day 28 post-challenge sample from one study participant that was previously not detected in our study. The participant was not symptomatic and presence of GII.4 was confirmed by RT-qPCR. A subsequent sample collected at day 35 post challenge was also tested by RT-qPCR did not contain the virus.

**Figure 2:**
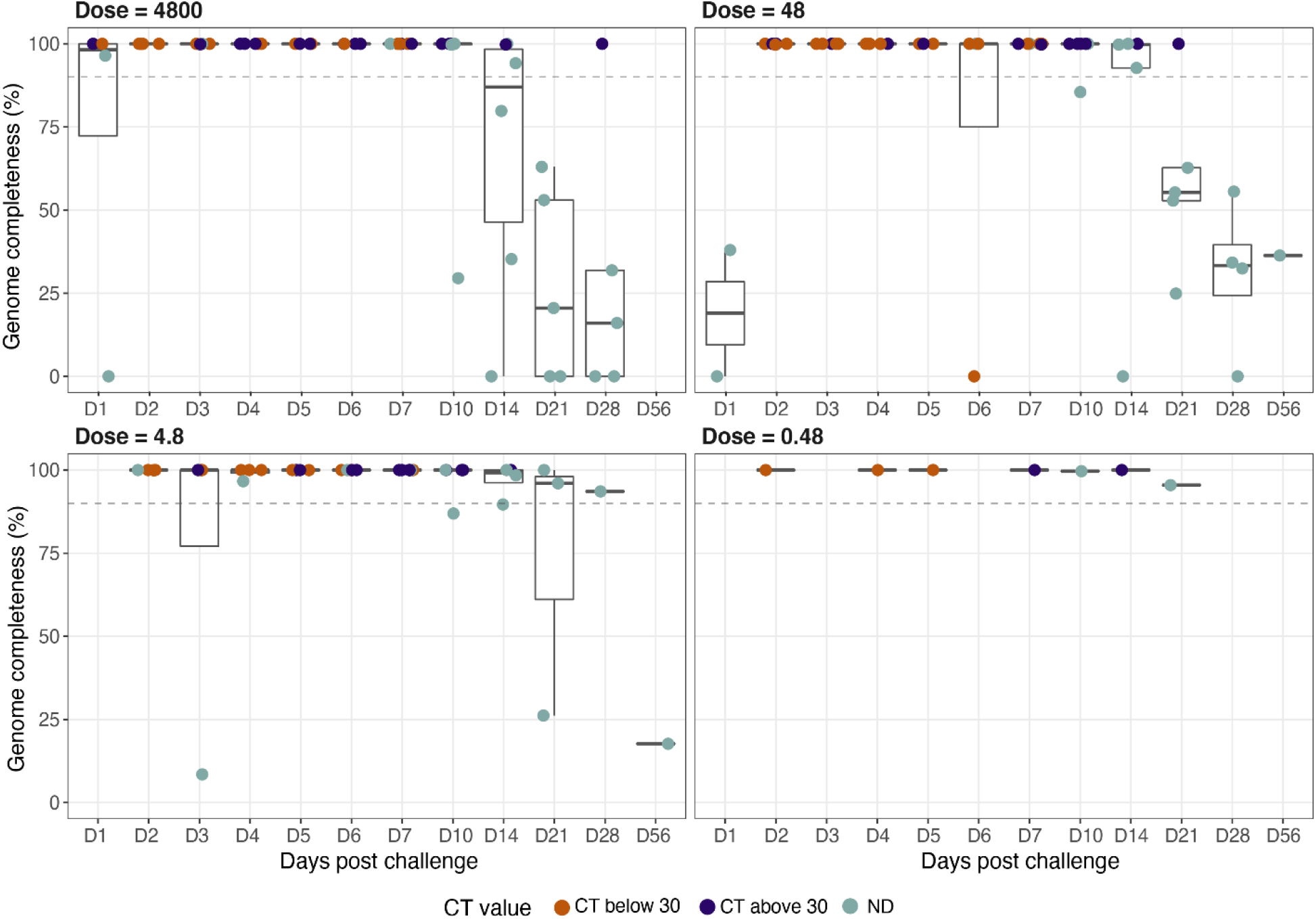
GI.1 genome assemblies in each challenge dose group. Box plots representing % genome completeness for study samples based on the four challenge dose groups (4800, 48, 4.8 and 0.48 RT-PCR units). Dotted line indicates >90% genome completeness. Days post challenge are indicated on the X-axis and % genome completeness is shown on the Y-axis. Samples are colored based on CT values post-library preparation. ND = not detected and refers to samples that were below the limit of detection by standard RT-qPCR assay.

**Figure 3:**
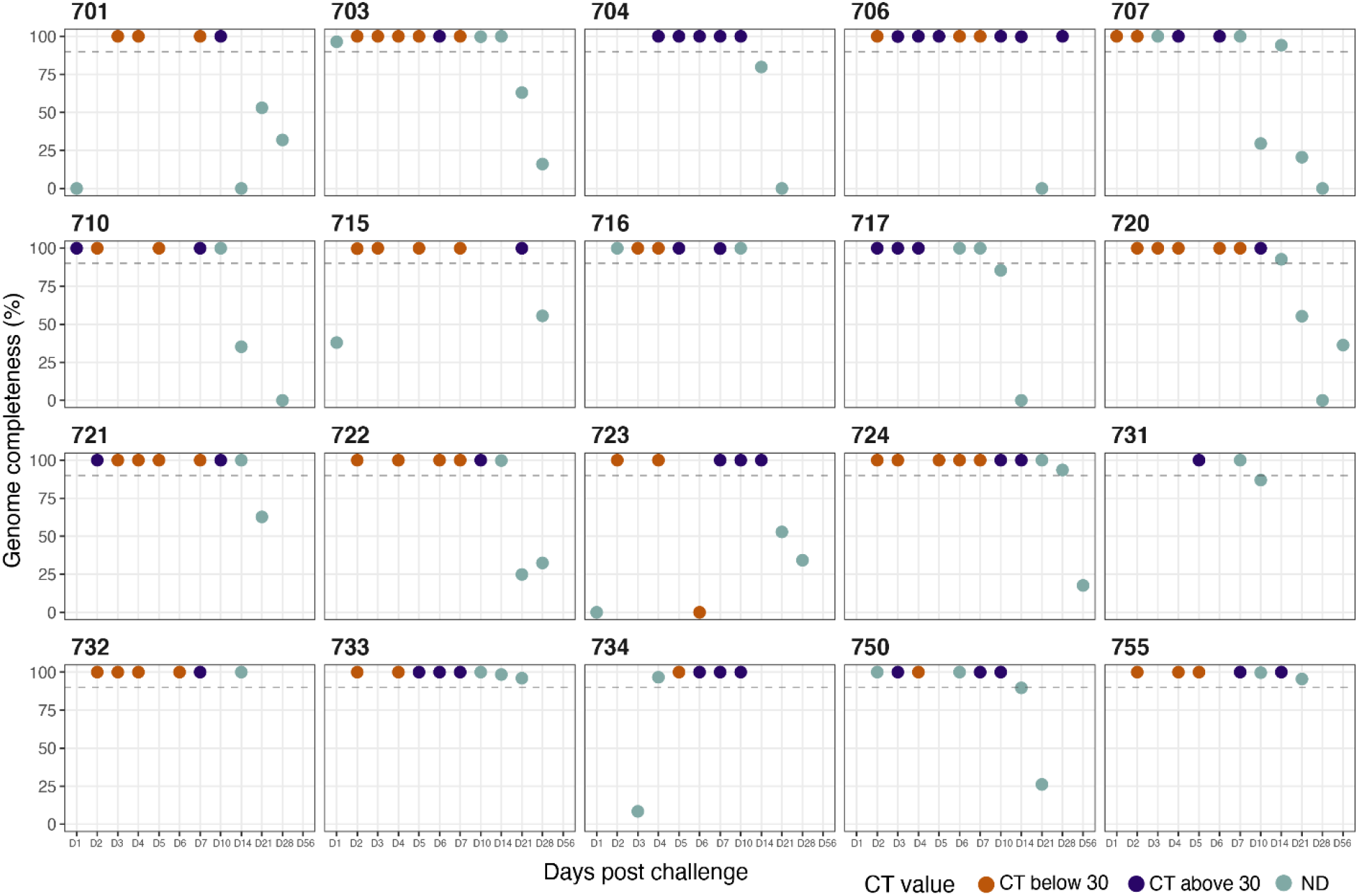
Genome assembly for each study participant infected with GI.1 norovirus. Samples with complete genomes (>90%, indicated by the dotted line) for study participant. Days post challenge are indicated on the X-axis and % genome completeness is shown on the Y-axis. Samples are colored based on CT values post-library preparation. ND = not detected and refers to samples that were below the limit of detection by standard RT-qPCR assay.

### Single nucleotide variant analysis shows limited conserved changes across persons

Analysis for single nucleotide variants (also referred to as alternate alleles) was performed on samples with complete genomes. The total number of mapped reads and frequency of detection of alternate alleles for all study participants is shown in **Figure 4A**. To avoid overinterpretation of results due to inherent errors in sequencing, we used a minimum of 20 reads per position and detection of alternate allele at > 10% frequency **(Fig 4B)** to identify single nucleotide variants. As shown in **Figure 5**, changes in the GI.1 genome were observed in all persons. However, except for the 3’ untranslated region, there were no conserved changes across study participants, irrespective of clinical outcome. Per person distribution of single nucleotide variants are shown in **Figure 6**. As seen from this analysis, many single nucleotide variants persisted from early time points in each person, suggesting limited conserved hotspots for immune pressure-driven changes in healthy persons.

**Figure 4:**
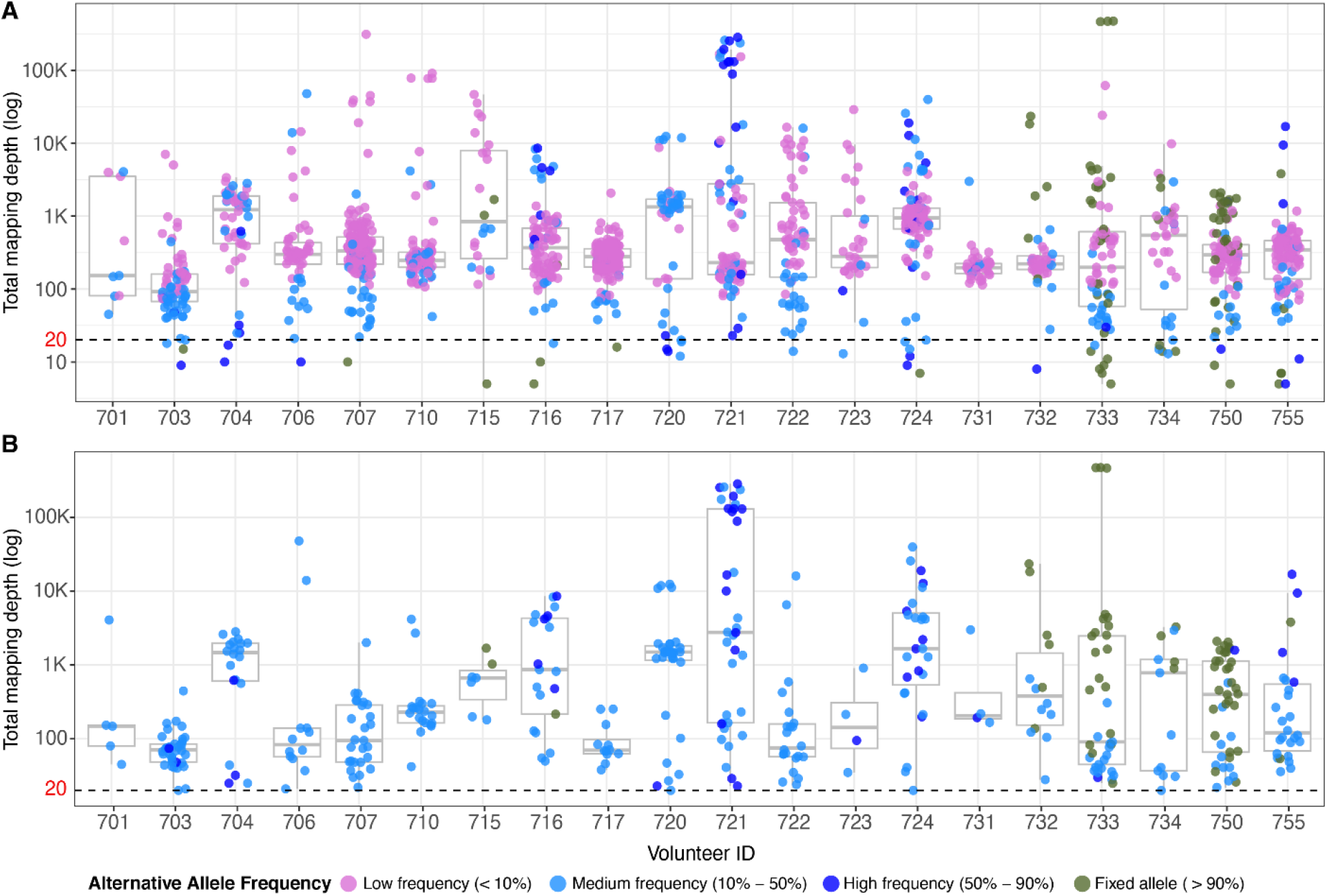
Detection of single nucleotide variants in study samples pre- and post-filtering. Detection of single nucleotide variants (alternate alleles) in samples from each study participant is shown prior to filtering (A) and post-filtering (B) where samples with less than 20 reads (below the dotted line) and low (<10%) allelic frequency (pink dots) were removed.

**Figure 5:**
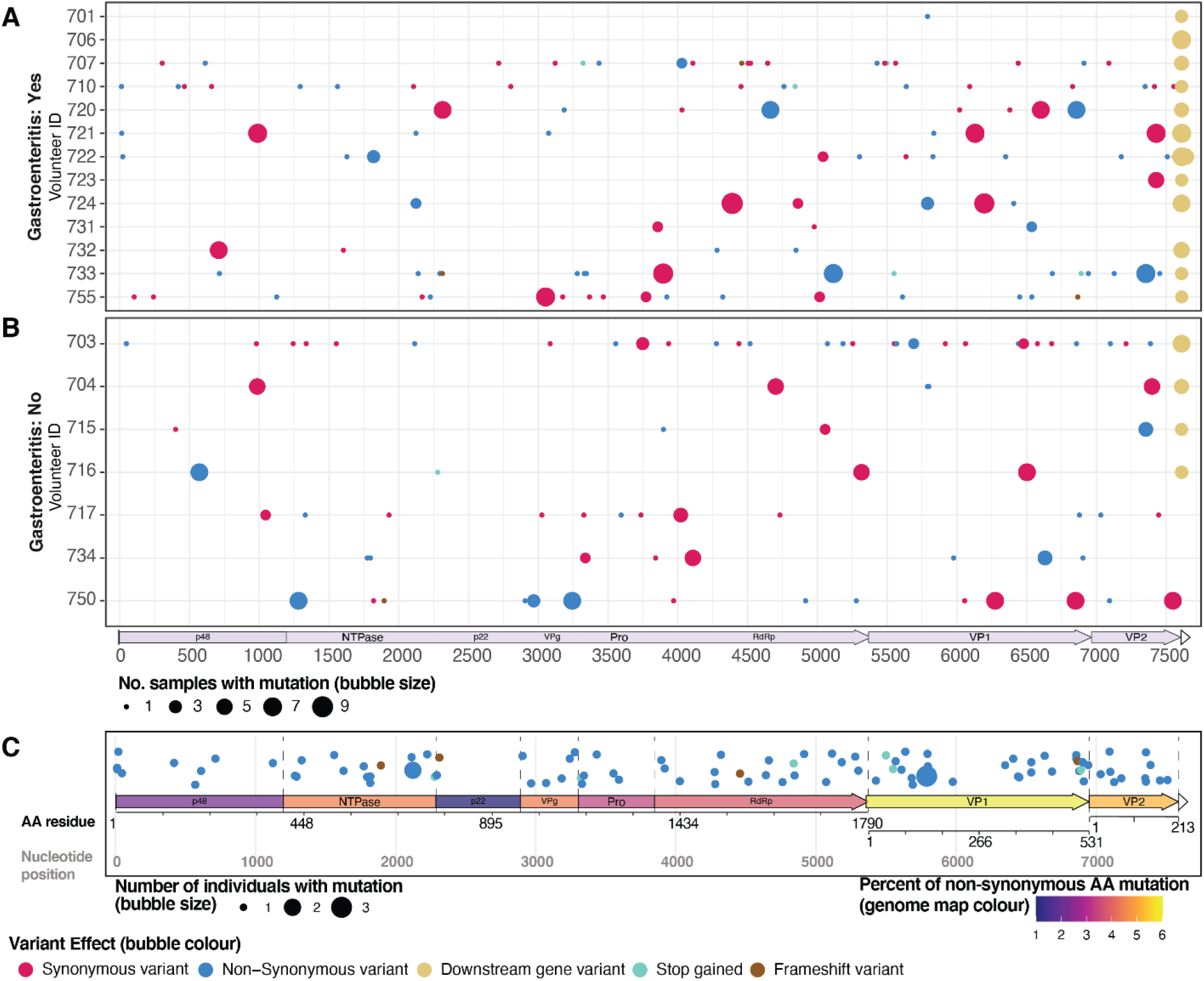
Amino acid changes due to single nucleotide variants in the GI.1 genome. Analysis of single nucleotide variants and corresponding amino acids changes in persons (A) with GI.1 associated gastroenteric and (B) those without clinical illness. The X-axis indicates different nucleotide positions of the GI.1 genome. The size of the bubbles is based on numbers of sample per person that show a change from the reference genome. (C) Positions where non-synonymous changes occur in the GI.1 genome are mapped for each protein with size of the bubble indicating number of persons with change from the reference genome and genome map colour indicating the % of non-synonymous changes in individual proteins.

**Figure 6:**
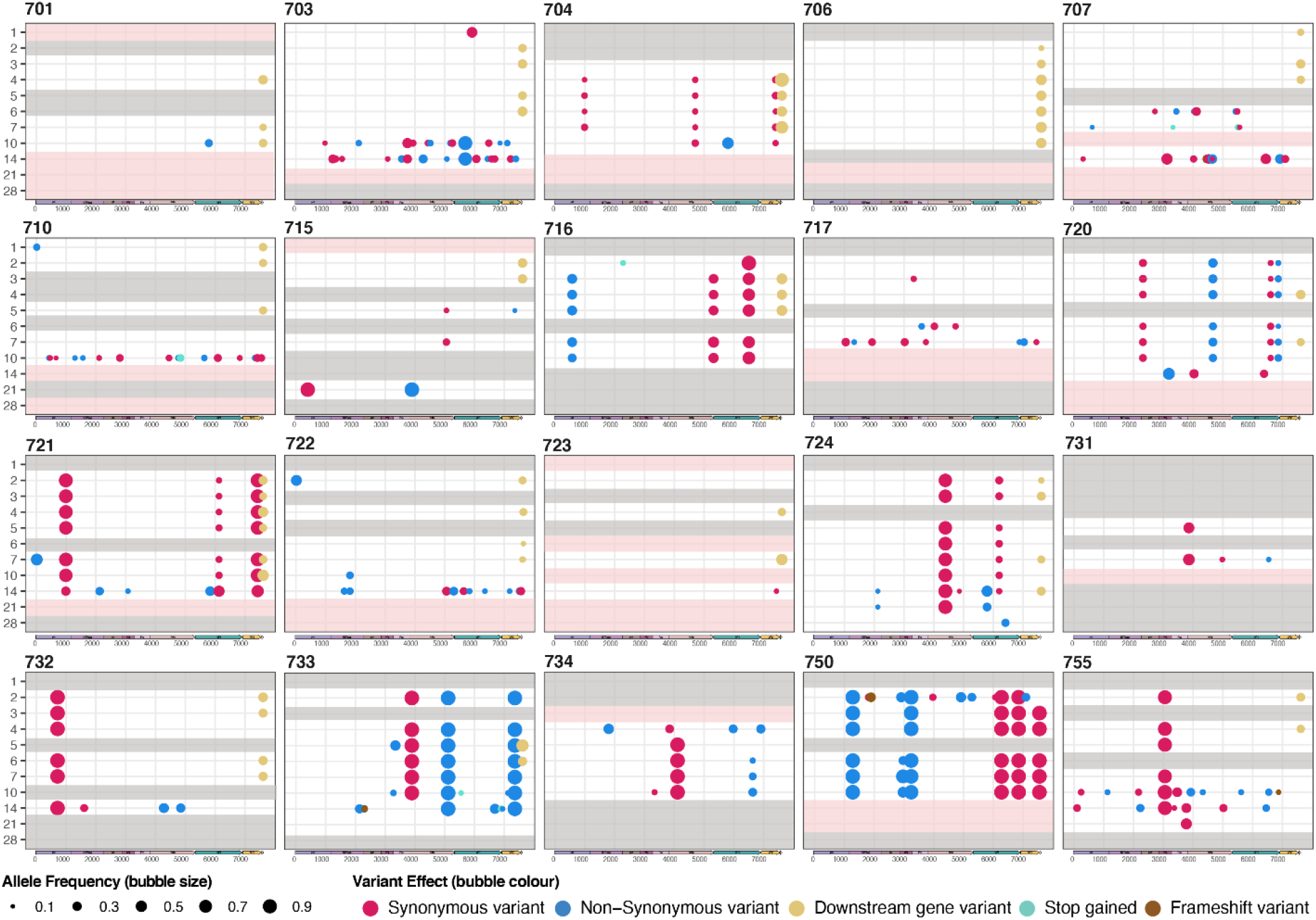
Single nucleotide variants in the GI.1 genome for each person in the study. Analysis of single nucleotide variants and corresponding amino acids changes for every study participant. Each row represents a day post-challenge with GI.1 virus. The size of the bubble indicates the frequency of detection of the alternate allele when compared to the reference. Data is shown only for time points where a complete genome was assembled. Gray lines across the panel indicates no sample while pink lines indicate incomplete genome.

### Non-synonymous amino acid changes are seen in structural and nonstructural proteins

Based on the single nucleotide changes, we next evaluated non-synonymous amino acid (aa) changes in each norovirus protein to determine if specific proteins were more subject to evolutionary pressure **(Figure 5C)**. Changes in VP1 were observed in 13/20 study participants (65%), although only three residues (aa 144, 393 and 500) changed in two or more persons **(Table 2)**. Although the number of study participants is small, we observed significantly higher number of changes in persons with GI.1 associated gastroenteritis than those without those without gastroenteritis (p = 0,01, chi-square test). Among non-structural proteins, the greatest number of changes were observed in the NTPase/NS3 (11/20 persons, 55%) and RNA-dependent RNA polymerase/NS7 (9/20 persons, 45%). A change from proline to serine at aa 308 was the only conserved change seen in 2 persons for NS7. Changes in the minor capsid protein were seen in 7/20 persons (35%). These data suggest similar levels of evolutionary pressure on non-structural proteins as seen with VP1.

**Table 2:** Numbers of amino acid changes from GI.1 reference genome for each protein. Number of unique positions showing a change from the reference genome and numbers of persons who showed any change from the reference is indicated.

| Protein | Positions with changes | % of persons |
| --- | --- | --- |
| NS1-2/p48 | 9 | 40% |
| NS3/NTPase | 14 | 55% |
| NS4/p22 | 2 | 5% |
| NS5/VPg | 6 | 20% |
| NS6/Pro | 6 | 20% |
| NS7/Pol | 19 | 45% |
| VP1 | 28 | 65% |
| VP2 | 10 | 35% |

## Discussion

What drives the emergence of new genotypes and variants has remained a long-standing question in the norovirus field given the remarkable genetic diversity of strains and the periodic emergence of new variants. Several different hypotheses have been proposed and tested including the potential for introduction of new viruses from immunocompromised persons with higher rates of virus evolution and zoonotic events leading to the acquisition of new ORF1 or ORF2 sequences into human viruses [11]. Increased risk of recombination in an immunologically naïve population such as children in day cares where there is a high norovirus disease burden is considered another source for emergence of new strains [24]. There is much less known about the evolution of human norovirus in healthy adults who are seropositive even though this population experiences substantial burden of infections and can shed virus in stool for extended periods of time even in the absence of clinical illness. In this study, we used capture-based sequencing to evaluate intra- and inter-host evolution of the prototype human norovirus strain, Norwalk virus in samples from a controlled human infection model. Our data showed changes in the GI.1 genome in all study participants but there was limited evidence for conserved changes between persons.

The accumulation of mutations in the antigenic epitopes of VP1 resulting in immune escape has been reported more frequently for GII.4 human noroviruses than other genotypes [30-33]. In that context, the lack of conserved changes in the GI.1 cohort may not be wholly unexpected. None of the changes seen in VP1 occurred in amino acids that bind histo-blood group antigens (HBGA) which serve as cellular attachment factors for human noroviruses [34]. However, of the three amino acid positions with changes conserved in two or more persons, position 500 is part of an epitope mapped by a nanobody that causes HBGA blockade activity through minor structural rearrangements of the protruding (P) domain of VP1 [35]. A change from asparagine to aspartic acid and tyrosine at position 393 was seen in two persons; this position is located within regions mapped by another nanobody that causes aggregation of virus like particles [36]. Complementary to our genomic analysis of stool samples, we recently used phage display of a GI.1 genomic library and deep sequencing to map antigenic epitopes in sequential serum samples from the controlled human infection study [37]. Antigenic landscapes varied between persons and new epitopes were seen 7 to 30 days post-challenge suggesting they the potential for their role in selection of variants. While the small sample size precludes any additional interpretation of the significance of these results, it would be important to analyze larger datasets, map changes to the structure of human noroviruses and evaluate their significance based on epitopes associated with virus neutralization and HBGA blocking.

Advances in sequencing methods and the potential to assemble complete genomes with high rigor offers the opportunity to gain new insights into the role of other proteins in norovirus emergence. Attention to the role of non-structural proteins in human norovirus evolution has largely focused on NS7 or the RNA dependent RNA polymerase (RdRp) [38]. This is primarily because the ORF1/ORF2 junction between NS7 and VP1 plays a major role in recombination and generation of chimeric viruses with potentially better fitness. Indeed, recombination with specific RNA-dependent, RNA polymerases have been associated with the emergence of GII.4 variants as well as other genotypes such as GII.2 and GII.17 [38]. While we observed changes in RdRp in our study, we also noted a high frequency of changes in NS3 (NTPase). Phage display of the GI.1 genomic library also resulted in the identification of 13 epitopes in nonstructural proteins samples, including 3 epitopes in NS3 [37]. NS3 is known to play an important role in norovirus replication including facilitating NS7-mediated RNA synthesis [39]. These data suggest NS3 may be an important protein to consider in the context of virus evolution.

The concept of non-GII.4 strains being more static compared to the globally dominant GII.4 strains has been described previously [23, 40]. Our results are consistent with this hypothesis although one limitation of the present study is that analysis was restricted to samples from the less prevalent GI.1 genotype. Another limitation is the small sample size; while we tested 156 samples in this study, these samples were collected from 20 infected persons limiting inter-host evolutionary analyses. Although we did not see differences in numbers of compete genomes based on challenge dose, whether there are differences in evolutionary patterns between persons with and without gastrointestinal symptoms is more difficult to conclude based on the small sample size. An increase in challenge dose was associated with more rapid shedding and symptom onset, and possibly increased severity in this cohort [41].

The capture probes-based sequencing method established in this work was highly sensitive and could be used to assemble complete genomes in samples with very low viral load and over extended periods of time. While the probe set was tested only on persons challenged with GI.1, the unexpected detection and complete genome assembly of a GII.4 Hunter variant suggests the potential application of this method for other human norovirus strains. This study provides proof of principle for using capture-based sequencing to study intra- and inter-host evolution of human noroviruses. Overlaying data on immune responses and antigenic epitopes with full length genome assemblies from individual study participants may provide new insights into intra-host evolution of human noroviruses.

## Acknowledgements

This work was supported by grants from the National Institutes of Health (U19 AI144297 and P01 AI057788).

